# Active perceptual learning involves motor exploration and adaptation of predictive sensory integration

**DOI:** 10.1101/2022.02.17.480969

**Authors:** Masato Hirano, Shinichi Furuya

**Affiliations:** Sony Computer Science Laboratories, Inc., Tokyo, Japan; NeuroPiano Institute, Kyoto, Japan

**Keywords:** Active perceptual learning, Active exploration, Predictive integration

## Abstract

We perceive the external world through both externally generated and self-generated sensory stimuli (i.e., passive and active perception). While the performance in passive perception is improved by training or repetition through the functional and structural reorganization of the central nervous system (i.e., perceptual learning), the mechanisms by which perceptual learning occurs in active perception remain unclear. Here, we sought to explore the mechanisms underlying the improvement of active somatosensory perception and sensorimotor skills through active perceptual learning. Because we previously found that active perceptual learning depends on the expertise of the motor task to be performed, this study focused primarily on trained individuals. To this end, active haptic training (AHT) that targets the active somatosensory perception during the piano keystroke was used as a means of inducing active perceptual learning. We found four main results. First, participants actively modulated the muscle coordination patterns to optimize the movements required by the task through AHT without any explicit instruction on it by the experimenter, suggesting the involvement of active exploration in active perceptual learning. Second, AHT increased the relative reliance on afferent sensory information relative to the predicted one during the piano keystroke. Third, perceptual sensitivity of externally generated keystroke motions remained unchanged through AHT. Finally, AHT improved feedback control of repetitive keystroke movements in expert pianists. These results suggest that active perceptual learning involves changes in both the predictive integration process and active exploration and that the improved feedback control of fine movements benefits from the improvement in active perception.

**Significant statement:** Our ability to perceive both externally generated and self-generated sensory stimuli can be enhanced through training, known as passive and active perceptual learning. Here, we sought to explore the mechanisms underlying active perceptual learning by using active haptic training (AHT), which has been demonstrated to enhance the somatosensory perception of a finger in a trained motor skill. First, AHT reorganized the muscular coordination during the piano keystroke. Second, AHT increased the relative reliance on afferent sensory information relative to predicted one, in contrast to no increment of overall perceptual sensitivity. Finally, AHT improved feedback movement control of keystrokes. These results suggest that active perceptual learning involves active exploration and adaptation of predictive sensory integration, which underlies the co-enhancement of active perception and feedback control of movements.

## Introduction

Accurate perception of the external world and one’s own physical state is crucial for the control and learning of dexterous motor skills, which form the foundation of cultural behaviors such as music and sports and of industrial development. Perception is traditionally divided into two categories(*1–3*): passive perception, which refers to the perception of sensory events that are externally generated, and active perception, which involves the perception of sensory consequences resulting from self-generated movements. Previous studies have demonstrated that passive perception can be enhanced through functional and structural reorganization in the nervous system, a phenomenon known as perceptual learning (*4–8*). However, the mechanisms of active perceptual learning remain poorly understood.

Active perception involves at least two processes not present in passive perception that can influence perception: active exploration and the integration of afferent sensory information with predicted sensory information. Active exploration involves interacting with the external world by controlling one’s own body parts (*9*). For example, we may change how to touch an object to collect information about the weight, curvature, surface texture, and/or size of the object being touched (*10, 11*). Previous studies also have shown that different movements can produce different perceptual experiences, even when interacting with the same object (*12, 13*). This suggests that active perceptual learning may occur by discovering an optimal movement strategy that conveys finely tuned sensory information through active exploration, such as “perceptual sweet spot”. The second process, the integration of afferent sensory information from sensory organs with predicted sensory information, occurs when the brain combines incoming sensory inputs with estimates of those inputs based on efference copies of motor commands (i.e. forward model) (*14–16*). This process of predictive integration can influence active perception by increasing sensitivity to unpredicted sensory information that occurs during movement or decreasing the effects of neural noise that disrupts sensory and sensorimotor processing (*17, 18*). Previous studies have revealed interindividual variation in this predictive integration process, which can change with age and disease (*19, 20*). Therefore, these two features are likely to be involved in active perceptual learning. However, no studies have yet investigated whether active exploration of novel movements and fine-tuning of the predictive integration process underlie the mechanisms of active perceptual learning. It is also unclear how active perceptual learning enhances not only active perception but also the control of fine movements. We recently demonstrated that active somatosensory perceptual learning reduces the force variability in a rhythmic movement that consists of a repetition of discrete force production movements. However, it remains unclear whether this results from enhanced movement-to-movement feedback control due to the improved active somatosensory processing through active perceptual learning or from improved fine control of the discrete movement. These contrasting possibilities predict opposite behavioral results that the training effect varies depending on the feedback-related gain of movements, which represents the extent that motor execution utilizes sensory feedback information obtained from past motor actions(*21, 22*).

Recent studies have demonstrated that active somatosensory perceptual learning occurs specifically in individuals proficient in the movements involved in the perceptual task (*23*). These skilled individuals have precise control over their movements (*24*), allowing them to creatively explore optimal movement strategies that provide finely tuned sensory information. In contrast, unskilled individuals exhibit large unintentional variability in their movements, preventing active exploration. Additionally, skilled individuals have more precise forward models of action relevant to a task than unskilled individuals(*25, 26*), suggesting that the predictive integration process during an active perception task functions specifically for skilled individuals. These can explain why active perceptual learning only occurs in skilled individuals and suggest that the mechanisms of active perceptual learning cannot be fully understood without considering skilled individuals. In this study, we examined how active perceptual learning affects active perception and fine motor control by assessing effects of active somatosensory perceptual learning in expert pianists. We found that both the active exploration of a novel movement pattern and a change in the predictive integration process in the nervous system occurred collaterally along with improvement in the active somatosensory perception of a piano keystroke movement through active perceptual learning in skilled pianists. We also discovered that these changes impact feedback control of fine keystroke movements in expert pianists.

## Results

Four experiments were conducted to investigate the mechanisms underlying the perceptual learning of active somatosensory perception (i.e., active perceptual learning) in skilled pianists. A total of 120 pianists and 10 musically untrained healthy individuals participated in the study. All pianists had received piano training at a conservatory or had received extensive private piano training under the guidance of a professional pianist or piano professor.

### Evidence of active motor exploration in active perceptual learning

To determine whether perceptual learning of active perception involves the active exploration of novel movements, we compared muscle activity patterns in keystroke movements before and after active haptic training (AHT) which was designed to enhance somatosensory discrimination perception of the weight of a piano key during a keystroke movement (*23*). In total, 24 pianists underwent an active perception test and the AHT in this experiment. In both tasks, they were instructed to strike a piano key attached to a haptic device twice in succession using their right index finger in each trial. The haptic device increased the key weight during either the first or second keystroke by pulling up the key. After performing two successive keystrokes, participants were asked to indicate which keystroke was perceived as heavier. In the active perception test, we used a Bayesian staircase procedure known as ZEST(*27*) to determine the discrimination threshold of the difference in the weight of the key (i.e., weight discrimination threshold: WDT). In the AHT, participants received feedback on whether their answer was correct or incorrect (i.e., performance success) to maximize perceptual learning immediately after each trial completion. The AHT consisted of 20 blocks, each comprising 20 trials. In the first block, the haptic device loaded the key to the amount corresponding to the WDT predetermined by the active perception test performed before the training. In this experiment, participants were divided into two groups based on whether they received feedback on the correctness of their answer during the AHT (FB group, n=12 (11 females, 23.6±2.6 y/o) or not (noFB group, n=12 (8 females, 21.6±2.3 y/o)). Since our previous study showed no improvement in the WDT without providing reinforcement signals during AHT (*23*), the noFB group is a control to ensure that changes in the dependent variables are due to improvement in active perception rather than task repetition.

Figure 1A shows the WDT obtained before and after the AHT in both groups. A linear mixed effects model (LME: fixed effects: group, time, and their interaction; random effect: participants) revealed a significant interaction between the time and group factors on the WDT (χ^2^(1)=5.15, p=0.02), indicating that the effect of AHT on the WDT differed between the groups. Specifically, the AHT improved the WDT in the FB group (t=2.50, p=0.02), but not in the noFB group (t=-1.02, p=0.33). This result is consistent with previous findings and suggests that active perceptual learning occurs only when feedback is provided during the training session.

**Figure 1.**
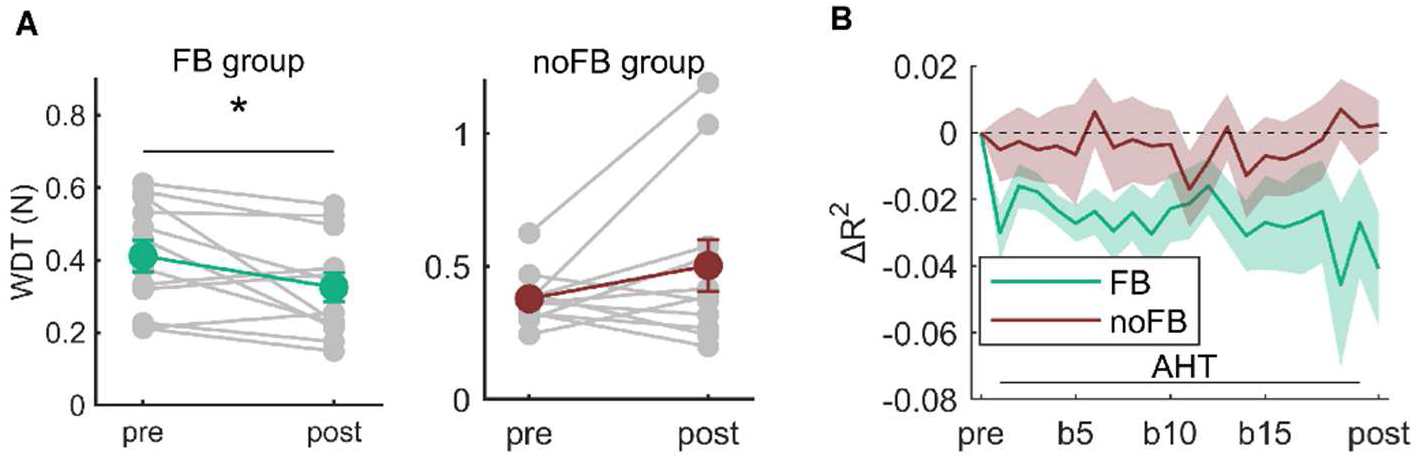
Change in muscular coordination through AHT. **A**: The average weight discrimination threshold (WDT) across participants obtained before and after AHT. **B**: The reconstruction quality of EMG signals of keystroke movement seen in the AHT and active somatosensory tests (in both pre- and post-tests) based on a linear summation of the two basis vectors extracted from the EMG signals in the active somatosensory test done before the AHT. Error bars and shaded areas represent the standard error. *: pre vs. post, p<0.05.

To determine whether each pianist acquired a novel movement that provides finely-tuned somatosensory information during keystrokes through AHT, we analyzed electromyography (EMG) signals from six muscles, that are involved in the index finger movements, obtained from the keystroke movements in both the active perception test and AHT. To quantify the changes in muscular coordination due to AHT, we used matrix factorization and data reconstruction techniques. First, we decomposed the muscle activities obtained from the active perception test done before the AHT into two distinct muscular coordination patterns (i.e., pre-patterns) and corresponding temporal coefficient patterns using a non-negative matrix factorization method (NMF) method. Then, we reconstructed the EMG signals of keystroke movements obtained during the AHT and from the active perception test done after the AHT according to the pre-patterns. If the muscular coordination patterns remain unchanged via AHT, the EMG signals in both the AHT and post-training test will span a similar low-dimensional subspace consisting of the pre-patterns. In this case, the quality of the reconstructed EMG signals, which is quantified by the coefficient of determination between the original and reconstructed signals, will be close to 1. The result showed that the quality of reconstructed EMG signals was smaller for the FB group than for the noFB group (Fig. 1B; LME, group: χ^2^(1)=3.98, p<0.05). In this experiment, the participants received no instruction regarding the effect of changing the motion of the keystroke on perception. Thus, the result suggests that pianists in the FB group actively explored movements by altering their muscular coordination patterns through the AHT. Whereas pianists in the noFB group did not reorganize the muscular coordination patterns.

### Increment of the relative reliance of afferent information on predictive information for active perception

The purpose of Experiment 2 was to determine whether the predictive integration process changes through AHT. To examine whether and how the integration of afferent and/or predictive information is modulated through AHT, we focused on sensorimotor attenuation. The amount of sensorimotor attenuation has been proposed as a measure of the relative reliance on afferent and predictive information at the integration of these two types of information (*18*). Therefore, assessing the amount of sensorimotor attenuation before and after AHT can provide insight into changes in the integration of these two types of information. The sensorimotor attenuation is typically assessed by a force-matching task where a participant is required to reproduce a perceived force by actively pushing a force sensor (*15*). We used a modified version of the force-matching task (i.e., keystroke force-matching task) in which participants were asked to reproduce target keystrokes that were passively generated by a haptic device as accurately as possible. We calculated the difference in the peak key-descending velocity between the reproduced and target keystroke as an index of the amount of sensory attenuation.

We first examined whether the keystroke force-matching task can successfully assess sensorimotor attenuation and whether the amount of sensorimotor attenuation differs between expert pianists and musically untrained individuals (nonmusicians). We hypothesized that pianists would show greater sensorimotor attenuation than nonmusicians because the reliance on predictive information derived from an internal model of keystroke movements in active somatosensory processing can be larger in pianists than in nonmusicians. Ten pianists (6 females, 22.9±3.1 y/o) and 10 nonmusicians (6 females, 23.4±3.4 y/o) participated in this study. They performed the keystroke-matching task. The haptic device passively moved the participants’ fingers by applying 4 different force levels (1.5 N, 2.1 N, 2.4 N, and 2.8 N) during the target keystrokes. Figure 2A shows the results of the keystroke force-matching task for participants in both groups. The pianists showed higher key-descending velocity of the reproduction strokes than that of the target keystrokes specifically in the low force levels (1.5 N and 2.1 N). This overcompensation is considered to result from sensorimotor attenuation and reflects the integration of the afferent and predictive information(*18*). In contrast, such overcompensation of the key-descending velocity in the reproduced keystrokes was not evident in nonmusicians. A LME (fixed effects: Group, Force, and their interaction; random effect: participant) yielded significant main effects of group (χ^2^(1)=5.97, p=0.01) and force level (χ^2^(3)=36.24, p<0.01) on sensorimotor attenuation (Figure 2B), which indicates that the keystroke force-matching task can successfully identify the sensorimotor attenuation in expert pianists when the haptic device moved the participant’s finger with a low level of force during the target keystrokes.

**Figure 2.**
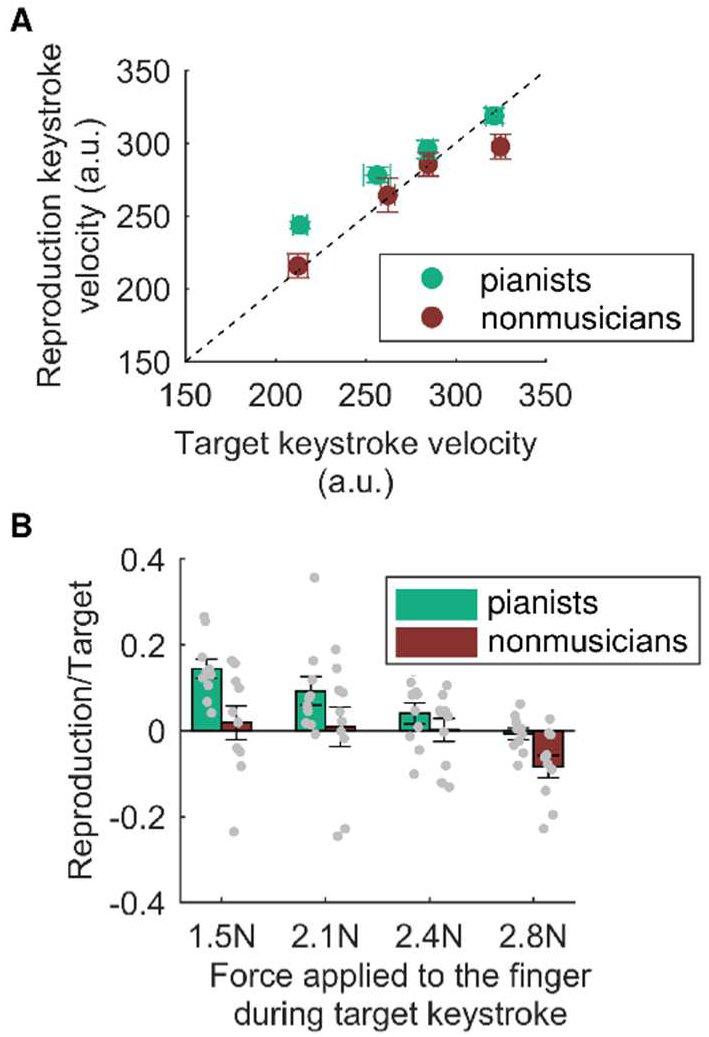
Keystroke force-matching task. **A**: Key-descending velocity generated by directly striking a piano key as a function of the externally generated target key-descending velocity. Green and brown plots represent the results obtained from the pianists and nonmusicians, respectively. **B**: The average values of velocity overcompensation across participants. A positive value indicates sensorimotor attenuation. Gray plots represent the individual data. Error bars represent the standard error.

We then examined the effect of AHT on sensorimotor attenuation. Thirty pianists were divided into two groups according to whether explicit feedback on performance success (i.e., reinforcement signals) was provided during AHT (FB (14 females, 23.6±6.1 y/o) and no-FB groups (15 females, 23.2±5.3 y/o), n=15 in both groups) and performed the keystroke force-matching task with a low level of force (1.5 N) before and after performing AHT. Figure 3A shows the changes in the WDT due to AHT in the FB and noFB groups. A LME (fixed effects: group, time, and their interaction; random effect: participants) detected a significant interactive effect between the group and time factors on the WDT (χ^2^(1)=5.18, p=0.02). Post hoc tests revealed a significant reduction in the WDT following AHT for participants in the FB group (t=4.66, p<0.01) but not for participants in the no-FB group (t=1.44, p=0.16). Furthermore, the WDT at the post-test differed significantly between participants in the two groups (t=-3.02, p<0.01), unlike the WDT at the pre-test (t=-0.51, p=0.62). Figure 3B shows the change in the amount of sensorimotor attenuation through AHT in participants in both groups. A LME (fixed effects: group, time, and their interaction; random effect: participants) yielded a significant interactive effect between the group and time factors on the amount of sensorimotor attenuation (χ^2^(1)=5.55, p=0.02). Post hoc tests revealed a significant reduction in the amount of sensorimotor attenuation through AHT in participants in the FB group (t=3.17, p<0.01) but not in participants in the no-FB group (t=-0.16, p=0.88). Because the amount of sensorimotor attenuation at the pre-test did not differ between the groups (t=0.93, p=0.35), these results suggest an increase in the relative reliance on the sensory afferent information to the predictive information through AHT.

**Figure 3.**
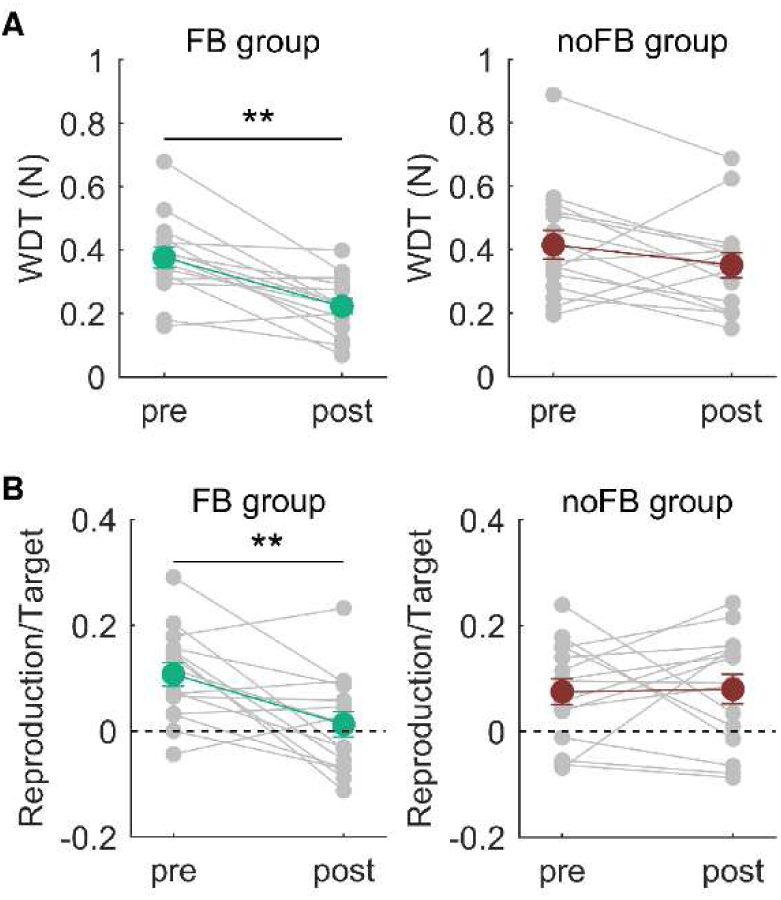
Reduction of sensory attenuation by AHT. **A**: The average weight discrimination threshold (WDT) across participants obtained before and after AHT. **B**: The average values of velocity overcompensation across participants in the active condition obtained before and after AHT. Error bars represent the standard error. **: pre vs. post, p<0.01.

### Passive perception unchanged through AHT

In this experiment, we examined whether the passive perception changed by AHT. Thirteen expert pianists (9 females, 20.2±1.6 y/o) participated in this experiment and performed the AHT with receiving feedback information(i.e., FB group). Before and after AHT, participants performed both active and passive perception tests. Specifically, for the active somatosensory test, we found a reduction in the WDT through AHT (pre-test: 0.56±0.21 N; post-test: 0.33±0.14 N (mean±SD); χ^2^(1)=19.90, p<0.01).

In the passive perception test, each participant was asked to discriminate the force of the two successive keystrokes done by the right index finger which are passively generated by the haptic device. The passive perception test consisted of 40 trials, and the force discrimination threshold (FDT) was calculated as the mean value of the posterior function obtained by the ZEST procedure across the 40 trials. We found no significant change in the FDT by AHT (pre-test: 0.20±0.08 N; post-test: 0.22±0.11 N (mean±SD); χ^2^(1)=0.13, p=0.72), which confirmed a lack of any change in passive somatosensory perception through AHT.

### AHT improved stroke-to-stroke feedback control of repetitive keystrokes

We previously found that the AHT reduces the force variability in repetitive keystrokes. This experiment further addressed the mechanisms underlying the effect of active perceptual learning in fine motor skills. To this end, we conducted a series of two experiments.

First, we tested whether the feedback-related gain of repetitive keystroke movements depends on the keystroke tempo. We hypothesized that at a faster tempo, the sensory feedback information obtained from a keystroke is more difficult to utilize in the force control of subsequent keystrokes (i.e., lower feedback-related gain) because the inter-keystroke interval and finger-key contact duration are too short to obtain task-relevant feedback information and to integrate it into subsequent motor outputs. To test this theory, 13 expert pianists (7 females, 22.9±1.9 y/o) performed a force-field adaptation task (Figure 4A) using repetitive keystrokes with the right index finger at various tempi (3, 2, 1, 0.5, and 0.25 Hz). In this task, participants were instructed to strike a force sensor attached to a piano key. They were asked to match the fingertip force applied to the key to a predetermined target force vector as accurately as possible while seeing the visual feedback on the error between the produced and target force vectors. In this task, the haptic device applied an external force to the finger in the direction of the abduction/adduction according to the perturbation schedule shown in Figure 4B (see details in the Methods section). The participants were required to compensate for this perturbing force. The averaged learning curves across the participants in each of the 5 tempo conditions are shown in Figure 4C. We calculated the deviation of the fingertip force from the target force as an error index. At the beginning of the adaptation phase, the error suddenly increased due to the perturbing force produced artificially by the haptic device. The error gradually decreased, although the slope of the decrement of the error appeared to differ among tempi values. To confirm this, we fitted the data with a single state-space model including both the feedback-related and retention gains. We found that the feedback-related gain increased as the tempo decreased (Figure 4D; A LME with a fixed effect of tempo and a random effect of participants: χ^2^(1)=21.71, p<0.01). In contrast, the retention gain did not differ across tempi (Figure 1E; χ^2^(1)=8.91, p=0.06). These results indicate that the feedback-related gain of the target keystroke task depends on the keystroke tempo.

**Figure 4.**
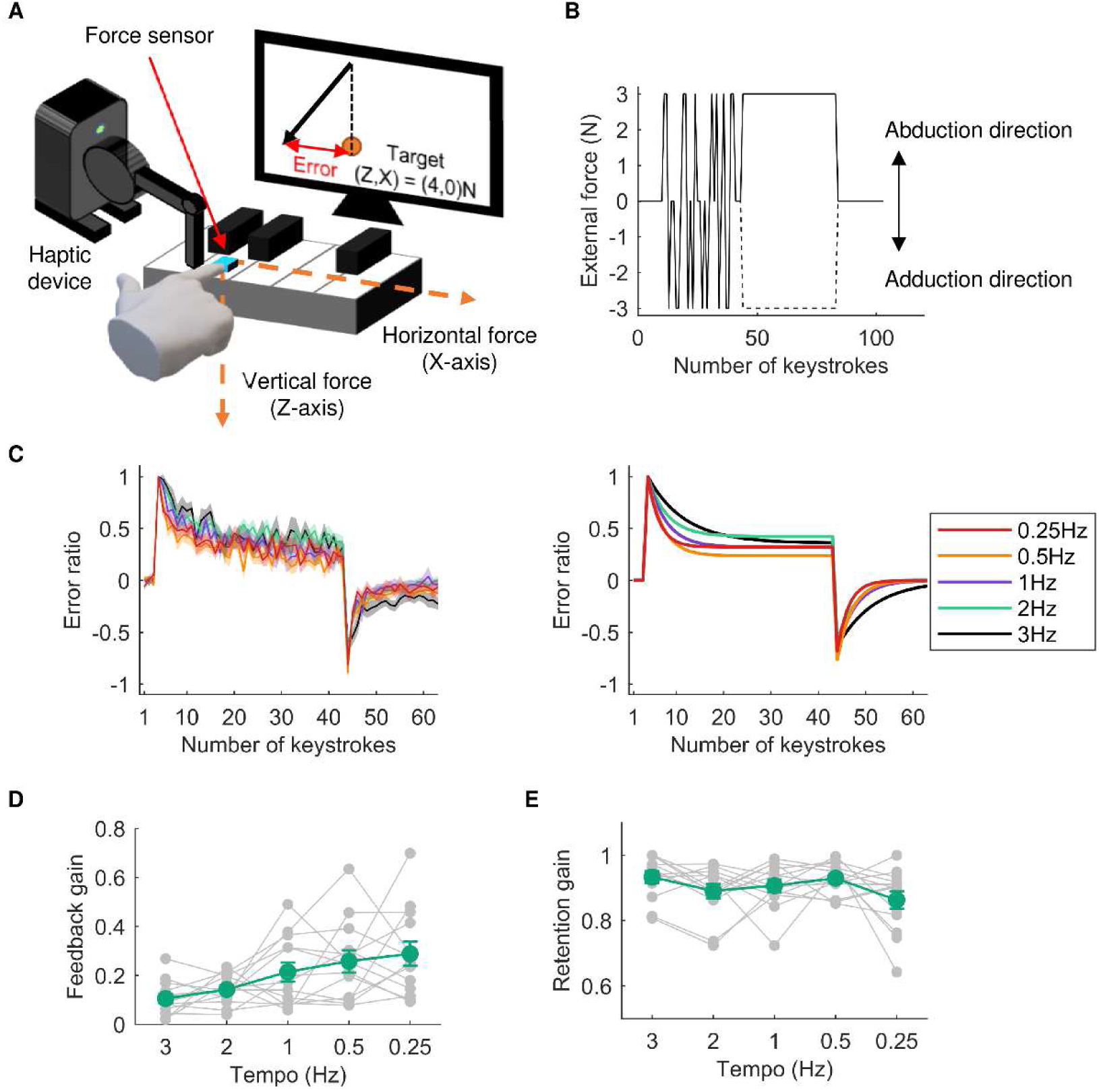
Force-field adaptation task in different tempi. **A**: An illustration that explains the force-field adaptation task. Participants were instructed to strike a piano key with predetermined vertical and horizontal forces at the fingertip of their right index fingers. A haptic device was attached to these fingers, and an external force was applied to these fingers along the horizontal direction during the random and adaptation phases. **B**: An example of the perturbation schedule during the force-field adaptation task. After the first 10 keystrokes, whether the external force applied to the finger and the direction of external force were determined randomly (i.e., random phase). After that, the external force was removed at 3-5 keystrokes (the number of keystrokes was randomly determined in each trial). During the subsequent 40 keystrokes, the external force was applied to the finger in either the abduction or adduction direction (i.e., adaptation phase). Finally, the external force was removed during the last 20 keystrokes (i.e., washout phase). **C**: The left panel shows average adaptation curves across participants in each tempo condition. The right panel shows the average adaptation curves fitted by the single state-space model. Each color line represents each condition, and the shaded areas represent the standard error. **D**, **E**: Average feedback-related gain (D) and retention gain (E) across participants in each tempo condition obtained by model fitting. The green and gray lines represent the mean and individual plots, respectively. Error bars represent the standard error.

We then examined the effect of AHT on the force variability of repetitive keystrokes performed at different tempi. Thirty-six pianists were categorized into two groups according to whether explicit feedback on performance success was provided during AHT (i.e., FB (14 females, 22.7±2.1 y/o) and no-FB groups (11 females, 22.8±2.0 y/o), n=18 in both groups). Before and after AHT, the participants performed the active perception test and a target keystroke task. Specifically for the active perception test, we found a significant reduction of the WDT through AHT in the FB group, but not in the noFB group (Fig. 5A and 5B: LME: interactive effect between the group and time factors, χ^2^(1)=4.19, p=0.04; pre-test vs post-test: FB group: t=3.51, p<0.01; noFB group: t=0.62, p=0.54).

**Figure 5.**
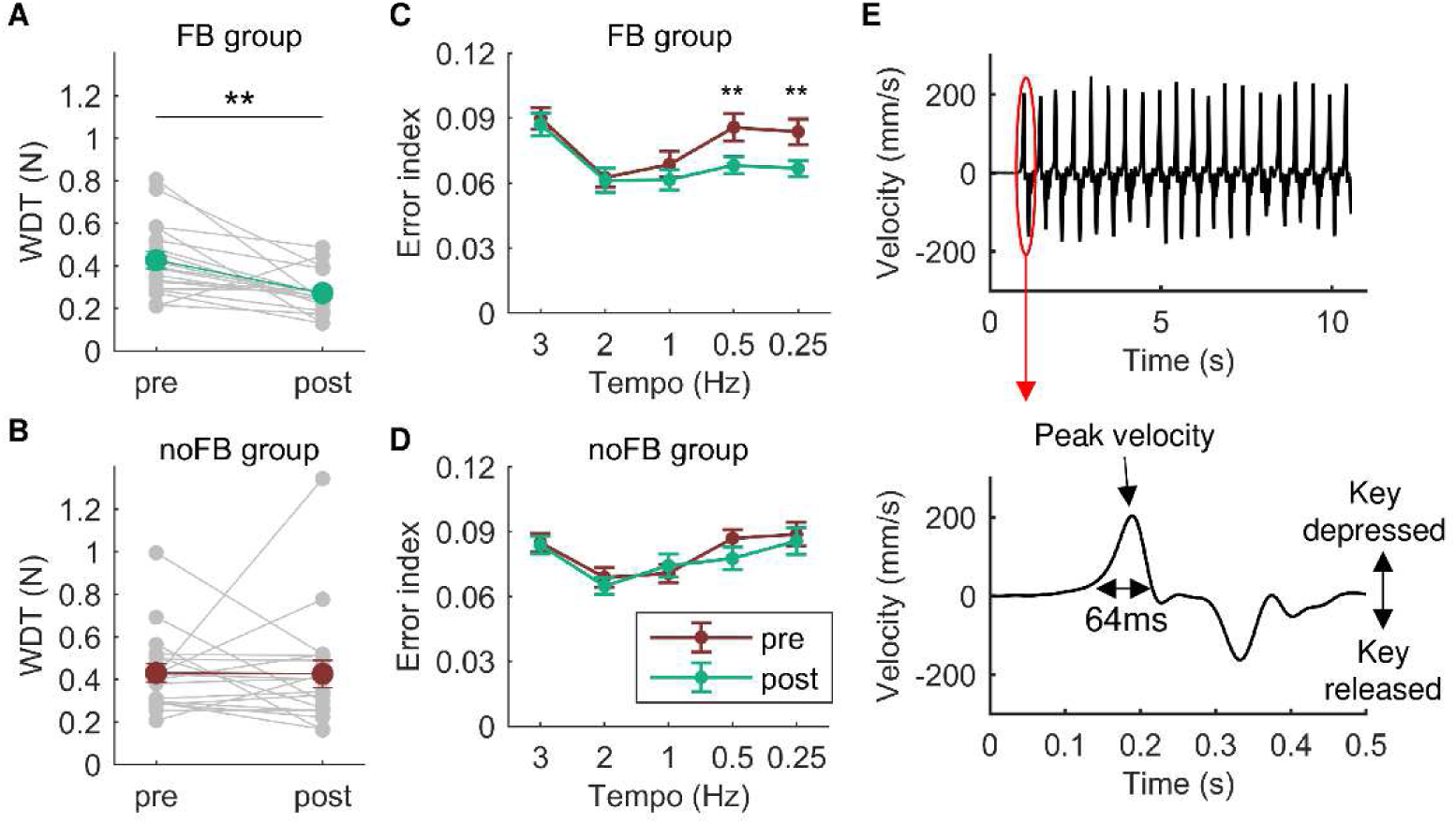
Effects of AHT on the performance of the repetitive keystroke task in different tempi. **A**, **B**: The average weight discrimination threshold across participants obtained before and after AHT in the FB (A) and noFB (B) groups. Green and brown lines represent the mean values across participants and the gray line represents the individual plots. **C**, **D**: The average error index across participants obtained before and after AHT for participants in the FB group (C) and no-FB (D) groups. **E**: An example of a time-course of key-movement velocity during the target keystroke task. The upper panel shows the whole time-course of the key-movement velocity in a single trial of the 2 Hz condition. The lower panel shows an enlarged view of the key-movement velocity in a single keystroke. The data show that the duration of the key-descending phase of a single keystroke was less than 100 ms. We calculated the coefficient of variation in the peak key-descending velocity across keystrokes as an error index. **: pre vs. post for participants in the FB group, p<0.01.

The participants performed the target keystroke task at each of various tempi before and after AHT. In this task, the participants repetitively struck a piano key 20 times at a rate of 3, 2, 1, 0.5, 0.25 Hz using the right index finger (Figure 5E). They were instructed to produce a predetermined key-descending velocity as accurately and consistently as possible. We calculated a coefficient of variation for the peak key-descending velocity values across the last 10 keystrokes as an index of the force production consistency (i.e., error index). Figures 5C and 5D show changes in the error index through AHT in both groups at each tempo. A LME on the error index (fixed effects: group, time, tempo, and their interactions; random effects: participant, participant*time, participant*Tempo) yielded significant group*time (χ^2^(1)=4.50, p<0.05) and tempo*time (χ^2^(4)=17.05, p<0.01) interactions. Simple effect tests revealed a significant reduction of the error index through AHT in the FB (t=-4.56, p<0.01), but not the noFB groups (t=-1.56, p=0.13). In addition, the error index was reduced only when performing the task at the slow tempi (0.25Hz: t=-3.55, p<0.01; 0.5Hz: t=-4.78, p<0.01; 1Hz: t=-0.31, p=0.75; 2Hz: t=-0.56, p=0.58; 3Hz: t=-0.65, p=0.52). In summary, AHT with the provision of FB information improved the target keystroke skill, and this effect was observed specifically at slow tempi with a high feedback-related gain of movements.

## Discussion

The present study explored the mechanisms underlying the enhancement of active somatosensory perception and sensorimotor skills through the AHT, a well-established method of inducing active perceptual learning in trained pianists but not in untrained individuals. We first found that active perception was improved and the muscular coordination for the keystroke movement was changed through AHT when the participants received reinforcement signals during the task. These results suggest that participants actively explored an optimal way of performing the keystroke movement to accurately perceive subtle differences in the weight of a piano key and that this active exploration depended on the presence of reinforcement signals. Second, we found that pianists exhibited stronger sensorimotor attenuation than nonmusicians, indicating a greater integration of predictive information into somatosensory perception during keystrokes in the former group. We also discovered that the amount of sensorimotor attenuation decreased through AHT, suggesting an increase in the relative reliance on afferent information relative to predictive information through AHT. Furthermore, the passive somatosensory perception was unchanged through AHT, indicating that AHT influences somatosensory functions responsible for active perception but not passive perception. Finally, found that the feedback-related gain of repetitive keystrokes depends on the movement tempi and that the effect of AHT on target keystroke skill was evident only when the task was performed at a slow tempo, during which the feedback-related gain of movements was high. These findings suggest that AHT improves stroke-to-stroke feedback control of repetitive keystrokes. Taken together, our results demonstrate that active perceptual learning involves both a change in the predictive integration process and active exploration and that the feedback control of fine movements benefits from the improvement of active perception.

### Active somatosensory perception

We replicated the previous finding that WDT in expert pianists decreased after AHT (*23*). The weight discrimination task that was employed in this study required perceiving a small difference in the weight of a key through keypresses based on somatosensory information derived from keystroke movements. While several studies have documented experience-dependent changes in passive somatosensory perception and related brain regions (*28–33*), it is largely unknown whether and how active somatosensory perception is enhanced through training. A key factor specific to active perceptual learning is the active exploration of optimal movements that provide enhanced task-relevant sensory information on a trial-to-trial basis. To investigate this possibility, we examined whether muscular coordination changed after AHT. To identify the change in muscular coordination, we used a series of multi-dimensional analyses. Since muscular coordination itself and how AHT changes the muscular coordination varied between individuals, univariate analyses, such as group mean of EMG signals across participants for each muscle, cannot fully capture the change in muscular coordination by AHT. Our multidimensional analyses, including matrix factorization and data reconstruction techniques, revealed that pianists who received reinforcement signals during AHT altered their muscular coordination, while pianists who did not have access to reinforcement signals maintained their muscular coordination throughout AHT. Importantly, the participants did not receive explicit instruction on the way of striking the key throughout the experiment, indicating that participants actively modulated their muscle activities. We propose that the participants were challenged to discover optimal keystroke movements that provided finely tuned somatosensory information based on the reinforcement signal, as active perception depends on movements (*12, 13*). For example, somatosensory afferents are likely to differ if the ways of touching keys vary between two keystrokes. Because the skin of the fingertip is deformable, deformations of the fingertip skin would vary depending on how to touch keys, which could alter the afferent inputs from the fingertip and the resulting perception(*34*). Thus, some keystroke movements may provide stronger task-relevant somatosensory information than others. The reinforcement signal drives active exploration of an optimal movement to improve discrimination performance in active perceptual learning. Furthermore, the unchanged passive perception during AHT in the presence of reinforcement signals suggests no changes in the sensitivity of the somatosensory system that is involved in passive perception. A theory of perceptual learning posits that reinforcement neural signals resulting from task performance enhance the processing of neuronal activity relevant to the task stimulus (*35–37*). The present study extends this theory by demonstrating that reinforcement signals influence how individuals interact with target objects/environments in active perceptual learning.

Another candidate process involved in active perceptual learning is the predictive integration of sensory information. Active perception is formed by integrating sensory information encoded in the afferent signals originating from the peripheral sensory receptors with predictive sensory information based on the internal forward models represented in the central nervous system (*38–40*). This predicts that changes in at least one of these two pieces of information underlie the improvement of the perceptual discrimination threshold in motion through AHT. We examined this prediction by probing sensorimotor attenuation, in which the sensory consequences of the voluntary action are perceived to be weaker than the same sensory event when it is externally generated(*18, 41*). The integration of predictive information with afferent information thus induces the attenuation of perceived sensory events. In Experiment 2, we found larger sensorimotor attenuation during keystrokes in expert pianists than in nonmusicians in low force levels, which means a larger reliance on sensory prediction for active somatosensory perception in the former individuals. Previous studies have demonstrated that sensorimotor attenuation is reduced when incoming afferent signals are unpredictable or temporally modulated (*42, 43*). Thus, in nonmusicians, a lack of accurate representation of novel mechanical characteristics of the piano may prevent sensory consequences of keystroke movements from being predicted. In addition, previous studies demonstrated modulation of sensorimotor attenuation in elderly individuals(*19*) and patients with schizophrenia(*20*). The present study extends these findings by providing the first evidence of expertise-dependent changes in sensorimotor attenuation. One may wonder why the sensory attenuation seen in the pianists appeared only at low force production. A previous study demonstrated that sensory attenuation occurs only when being tested with low, but not high force application in the force-matching task(*44*). Another study reported that the relationship between subjective force perception and applied force is not linear, rather subjective force perception is saturated at high force levels(*18*). Such a ceiling effect of subjective force perception could lead to an underestimation of the force to be applied. Thus, the sensory attenuation might be masked by the underestimation of high force due to the ceiling effect of subjective force perception. We further found the reduction of sensorimotor attenuation through AHT. This indicates an increase in the relative reliance on afferent sensory information for active somatosensory perception through AHT. The training effect can be mediated by functional changes in the peripheral system, such as enhancement of the sensitivity of fingertip somatosensory receptors. However, this is unlikely because the passive somatosensory perception was unchanged through AHT, in contrast to the enhancement of active somatosensory perception. Another possible reason for this outcome is a change in the movement that influences incoming sensory afferent signals. As described above, the muscular coordination patterns of the keystroke movement were reorganized through AHT. One plausible explanation for this is that pianists learned optimal keystroke strategy, which provides the highest proprioceptive sensitivity (e.g., striking the key with the most sensitive skin area of the fingertip, as has been argued as active exploration), through AHT. This enables pianists to obtain enhanced afferent signals during keystrokes, which increases the relative reliance on afferent signals over predictive signals. Together, the present results indicate that the pianists learned an optimal movement strategy that provided fine-tuned somatosensory information on the key weight during the keystrokes through AHT.

### Sensorimotor skill

We recently demonstrated that AHT improved the performance of the target keystroke task in expert pianists beyond the limit of performance improvement that can be achieved by conventional training (i.e., a mere repetition of task performance)(*23*). The target keystroke skill is characterized as repetitions of a single rapid keystroke movement, which suggests that both feedforward and feedback control processes are likely to contribute to the performance of this skill. Specifically, every single key depression is controlled mainly by the feedforward process because the duration of the movement is shorter than the sensory feedback delay (i.e., less than 100 ms) (see Figure 4A). This raises the possibility that refinement of the feedforward control of keystroke movements through AHT improved the target keystroke skill. Alternatively, a stroke-to-stroke feedback control process is an alternative possibility underlying the improvement of the target keystroke skill through AHT since somatosensory information obtained from a keystroke can be utilized to adjust the subsequent keystrokes. The present study tested which process improved through active perceptual learning. First, we found that the feedback-related gain was larger at slower keystroke tempi with longer duration to integrate the feedback somatosensory information obtained from a keystroke into motor outputs of the subsequent keystroke (i.e., sensorimotor feedback loop). Then, we demonstrated that AHT improved the target keystroke skill performed only at slow tempi, which indicates that AHT improves the sensorimotor feedback control of repetitive keystroke movements. AHT involves repeatedly discriminating unexpected components (i.e., increment of the weight of the piano key) based on the somatosensory information derived from one’s movements. Because movement control and learning rely heavily on corrective mechanisms based on errors between perceived and expected motor actions (*45, 46*), it is possible that AHT enhanced the somatosensory-based error-detection function and thereby improved the stroke-to-stroke feedback control component of the target keystroke skill.

### Limitation

There are at least two potential limitations in this study. One is that the same participant performed the force-field adaptation task multiple times in Experiment 4-1 (i.e., 5 tempo conditions × 2 times = 10 times), which may have elicited a saving effect. Saving is the ability of past learning to enhance subsequent learning(*22, 47–49*). Thus, it is possible that the adaptation rate might differ between data measured at the beginning and at the end of the experiment. To minimize this possibility, we randomized the order of tempo conditions across participants. We also included a random phase in which the external force was applied randomly in the direction of abduction or adduction. Previous studies demonstrated that exposure to two opposite perturbations during movements inhibits adaptation learning of both types of perturbations, which interferes with the formation of motor memories(*50, 51*). The random phase was always followed by the adaptation phase to prevent the saving effect. Furthermore, to prevent prediction of the direction of external force during the adaptation phase, the external force was applied once in the abduction direction and once in the adduction direction at each tempo in random order. These schemes might minimize the suppression of the potential effects of saving on the results of the adaptation task.

Another limitation is that while AHT improved the target keystroke skill performed at a tempo of 2 Hz in the previous study(*23*), the same training did not improve the skills of participants in the present study. A contrasting difference is that participants performed the target keystroke task using the right ring and index finger in the previous and present study, respectively. The ring finger has innately lower finger independence and controllability of movements compared with those of the index finger(*52, 53*), which is confirmed by a superior performance index of the target keystroke skill in the present study over those observed in the previous study. This suggests a potential for improvement of feedforward motor control specifically in the ring finger. Thus, it is possible that, in the previous study, AHT improved both the feedforward and feedback motor control of the right ring finger in expert pianists, but AHT only improved the feedback motor control of the right index finger of participants in the present study.

## Conclusion

The present study investigated the mechanisms underlying active perceptual learning by focusing on the effects of AHT on active somatosensory perception and fine motor control of piano keystrokes in pianists. A series of four experiments provided evidence that AHT led to the active exploration of optimal movements that provide fine-tuned somatosensory information and that AHT increased reliance on the somatosensory afferent information derived from one’s own movements in the somatosensory process in motion. These mechanisms can underly active perceptual learning and optimize feedback sensorimotor control.

### STAR Methods Participants

In total, 120 healthy pianists and 10 nonmusicians participated in this study. All of the pianists majored in piano performance at a musical conservatory and/or had extensive and continuous private piano training under the supervision of professional pianists and/or piano professors. All participants gave their written informed consent before engaging in the experiments. All experimental procedures were carried out in accordance with the Declaration of Helsinki and were approved by the ethics committees of the Sony Corporation. To remove the effects of auditory feedback on the performance of all experimental tasks, we muted a piano and instructed each participant to perform each behavioral task while listening to white noise sounded from headphones worn on their ears.

### Recording of piano key movements

The vertical motion of the piano key was measured using an optical distance sensor placed under the key at a sampling rate of 500 Hz. The sensor values were low-pass filtered (fifth-order zero-lag Butterworth filter, cutoff frequency: 24 Hz) and differentiated to obtain the velocity values.

### Recording of electromyogram activity

Surface electromyogram (EMG) activities were recorded from the finger intrinsic (i.e., first and second dorsal interosseous muscles (1DI and 2DI)) and extrinsic muscles (i.e., distal and proximal part of flexor digitorum superficialis muscle (dFDS and pFDS), extensor digitorum muscle (ED), and extensor indicis muscle (EI)) of the right hand with Ag/AgCl surface electrodes (Trigno TM mini, Delsys, USA). The analog signals were digitized at a sampling rate of 1 kHz and then fed into a computer using a custom-made program for offline analyses (LabVIEW, National Instruments). The raw EMG signals were high-pass filtered at 50 Hz using a zero-phase-lag sixth-order Butterworth filter and were demeaned, digitally rectified and low-pass filtered at 20 Hz (a zero-phase-lag sixth-order Butterworth filter). Finally, the EMG signals of each muscle were integrated over a 5 ms window and divided by the maximum EMG obtained during a maximum isometric contraction task done prior to the experiment. During the isometric contraction task, each participant was instructed to produce a maximum force of the index finger abduction, adduction, extension, and flexion while the finger joint was immobilized by the participant’s left hand.

### Active somatosensory test

The active somatosensory test was designed based on our previous study (*23*). It involved a weight discrimination task that assessed the weight discrimination threshold (WDT) as to changes in the key weight when striking the piano key with the right index finger. Participants were instructed to strike a key at a peak key-descending velocity amounting to 50% of their maximum (i.e., maximum peak key-descending velocity: MVK) twice in succession at each trial. A warning message of “too weak” or “too strong” was displayed on a monitor placed in front of the participants when the peak key-descending velocity was outside 35-65% of the MVK. During only one of the two strikes, a haptic device (Geomagic Touch X, 3D Systems), attached to the key by a double-sided tape, increased the key weight by pulling up on the key. Subsequently, participants answered which keystroke they perceived as heavier. The active somatosensory test consisted of 40 trials, and the WDT was calculated by a Bayesian staircase procedure (i.e., ZEST)(*27*) through the 40 trials. Briefly, ZEST is an efficient method of measuring thresholds of discrimination perception. We first specified prior knowledge, namely, a guess (=-0.40N in a logarithmic (log) scale) and an associated standard deviation (=2N in a log scale) for the probability of threshold values. These values were used to calculate the initial probability density function (pdf) of the threshold by assuming it to be a Gaussian function. The pdf was updated every trial based on the result of the discrimination test. The stimulus intensity and the final estimate of the threshold were chosen to correspond to the mean of the latest pdf. Because the ZEST worked based on a log scale, log-scale values were used for statistical testing. By contrast, for visualization, we adopted linear scale values to increase the understandability.

### Active haptic training

Active haptic training (AHT) was designed based on our previous study (*23*). The general procedure was the same as the active somatosensory test. But, following each trial in the AHT, we provided visual feedback on whether the answer was correct or incorrect (i.e., a reinforcement signal on performance success). The AHT consisted of 20 blocks, each of which had 20 trials. In the first block, the haptic device loaded the key to the amount corresponding to one’s WDT predetermined by the active somatosensory test performed prior to the training. To maintain task difficulty and facilitate perceptual learning during the training session, the load for the next block was reduced by 10% from the load for the previous block when participants correctly answered more than 80% of trials within a block. In experiments 1, 2-2, and 4-2, half participants performed this task with the provision of visual feedback on the correctness of the answer (i.e., FB group) whereas the other half participants did not receive the visual feedback as a control group (i.e., noFB group).

### Experiment 1

The purpose of experiment 1 was to determine whether perceptual learning of active perception involves the active exploration of novel movements in skilled pianists. To this aim, we recorded muscle activity while participants performed the active somatosensory test and compared the muscle coordination during striking a key before and after the AHT.

### Protocol

Twenty-four pianists participated in this experiment and were divided into two groups (i.e., FB, and noFB groups, both n=12). Because one pianist in the noFB group had an abnormal WDT (greater than 1.3N), we excluded this pianist’s data from the analyses. Before and after the AHT, the participants performed the active somatosensory test (i.e., pre-test and post-test). During the active somatosensory tests and the AHT, we recorded the EMG signals from the six muscles while striking a key. The EMG signals were segmented in epochs from -200 ms after the onset of the key press to 200 ms after the onset of the key release. The onset of both the key press and the key release was identified from the MIDI events generated by the electric piano, which were synchronized with the EMG recordings. Because the weight discrimination task included in both the active somatosensory test and the AHT required each participant to strike a key twice in each trial, we obtained two sets of EMG signals in a single trial. During one of these two keystrokes, the haptic device loaded the key weight. However, the amount of loading varied between trials and participants, so we only used the data from the keystroke during which the haptic device did not load the key weight for offline analyses.

To identify changes in muscular coordination through AHT, we performed a series of multidimensional analyses. First, we decomposed the preprocessed EMG signals obtained from the pre-test into multiple sets of a muscular coordination pattern and a corresponding temporal coefficient pattern using a non-negative matrix factorization (NMF) analysis. We calculated the variance accounted for (VAF) to determine to the extent to which the input EMG data could be reconstructed from the decomposed data. The VAF is defined as the ratio of the variance between the residuals of the reconstructed and input data (i.e., VAF is equal to the coefficient of determination). We decomposed the EMG signals into two muscular coordination patterns. This is because a keystroke consists of a key press and release phase and most of the data variance in the EMG signals (>80%) spans two dimensions in most participants. Since NMF involves an iterative algorithm and the initial weight values of the iteration are determined randomly, we first repeated the NMF decomposition 20 times and used the solution that gave the highest VAF in the subsequent NMF as the initial weight values to increase the robustness of the decomposition performance.

After performing the NMF, the preprocessed EMG signals obtained from the post-test and the AHT were reconstructed based on the muscle coordination patterns extracted from the pre-test and optimal temporal coefficient patterns. The optimal temporal coefficient patterns were identified by optimizing the temporal coefficient patterns while maintaining the muscular coordination patterns, in order to minimize the error between the original and reconstructed EMG signals. If a given set of EMG signals spans in subspaces similar to those obtained in the pre-test, the quality of the reconstructed signals quantified by the VAF will be close to 1, whereas it will be close to 0 if the muscular coordination differs between two data sets.

### Experiment 2-1

The purpose of this experiment was to examine sensorimotor attenuation during keystroke movements between pianists and nonmusicians. Ten pianists and 10 nonmusicians performed the keystroke-matching task.

### Keystroke force-matching task

As a robust quantitative measure of sensorimotor attenuation, the force-matching task has been used in previous studies, in which participants were asked to produce the force that matched the target force produced by a torque motor (*15*). We customized this task for piano keystrokes. In this task, participants were asked to reproduce a target keystroke movement that was passively generated by the haptic device. The haptic device applied downward force to the participants’ right index fingers with a predetermined force level (1.5, 2.1, 2.4, and 2.8 N) at a tempo of 2 Hz. The force was randomly selected across trials and was applied to the finger 10 times in each trial, and each applied force lasted 200 ms. Following passive movements, the haptic device was automatically detached from the finger, and the participants were instructed to reproduce the target keystroke generated by the haptic device as accurately as possible (active condition). In a control condition that evaluates haptic pressure perception, participants controlled keystroke movements indirectly through the haptic device at the reproduction session. The participants can manipulate the force level of the haptic device via piano keys using their left hands. Each of the 5 keys corresponded to each of the following functions: ‘increase force function’, ‘decrease force function’, ‘reproduction function’, ‘target function’, and ‘finish function’. Participants can manipulate the force level in the step of 0.1 N by selecting the ‘increase/decrease force function’ for the passive keystrokes triggered by the ‘reproduce function’. In each trial, participants can confirm the target keystrokes and reproduction keystrokes any number of times by pressing the keys corresponding to ‘target function’ and ‘reproduction function’, respectively. During the control condition, participants can see the PC monitor where the correspondence table between the keys and functions is displayed. We instructed each participant to close his or her eyes when pressing the keys corresponding to the target and reproduction functions and to match the peak key-descending velocities of the target and reproduction keystrokes. Once they perceived that the two types of keystrokes were the same, they were asked to press the ‘finish function’ to proceed with the task.

We calculated the ratio of a difference in the peak key-descending velocity between the target keystroke and reproduced keystroke as an index of the amount of sensory attenuation. The positive and negative values indicate stronger and weaker keystrokes in the reproduced keystroke than in the target keystroke, respectively.

### Experiment 2-2

The purpose of this experiment was to examine the effect of AHT on sensorimotor attenuation. Thirty pianists participated in this study.

### Protocol

The pianists were categorized into two groups. One group of participants can receive feedback information about whether their answers were correct during AHT (FB group, n=15). The pianists in the other group (no-FB group, n=15) cannot receive such feedback information during AHT. Before and after AHT, the participants performed the active somatosensory test and the keystroke force-matching task with the active condition. Because the sensorimotor attenuation was evident only in low-force levels, the participants performed the keystroke force-matching task only with a low-force level (1.5 N).

### Experiment 3

This experiment examined whether AHT modulates passive somatosensory functions. Thirteen pianists performed a passive somatosensory test before and after AHT. All participants received the visual FB on the correctness of the answer during the AHT (i.e., FB group).

### Passive somatosensory test

The passive somatosensory test involved a keystroke strength discrimination task that assessed the passive discrimination threshold regarding changes in the strength of keystrokes generated by the haptic device attached to the participants’ right index fingers. The haptic device applied a downward force to the finger to strike a key twice in succession at each trial. The strength differs between the two passive keystrokes. Subsequently, participants indicated which passive keystroke they perceived as being more difficult. The passive somatosensory test consists of 40 trials, and the FDT was calculated by the ZEST.

### Experiment 4

Experiment 4 was designed to clarify the mechanisms underlying the effect of AHT on the participants’ target keystroke skills. We conducted two experiments to examine the hypothesis that AHT improves feedback motor control.

### Experiment 4-1

The purpose of this experiment was to examine whether the feedback-related gain of repetitive keystroke movements depended on the keystroke tempo. Thirteen pianists performed the force-field adaptation task.

### Force-field adaptation task

The participants performed repetitive strikes of a piano key at one of the five tempi (3, 2, 1, 0.5, and 0.25 Hz) using the right index finger to which the haptic device was attached. In this task, participants were instructed to strike the piano key so that a fingertip force applied to the key could match a predetermined target force vector. We recorded the 2-dimensional fingertip force applied to the key during each keystroke using a force sensor (USL06-H5-100N. Tech Gihan, Japan) attached to the key. We calculated the two-dimensional fingertip force vector (i.e., the X and Z axes, each of which corresponds to the finger abduction/adduction and downward directions, respectively) at the moment when the Z axis force reached a maximum value during each keystroke. The pianists can see the force vector and a target force vector for every keystroke, which were displayed on a PC monitor located in front of them. We instructed pianists to match the fingertip force vector to a target force vector ((X, Z) = (0N, 4N)) as accurately as possible for each keystroke. During the first 10 keystrokes, the haptic device did not apply any force to the participants’ fingers. After these 10 keystrokes, the haptic device applied an artificial force to the participants’ right index fingers in the direction of abduction or adduction (i.e., X axis). The artificial force direction and whether the artificial force was applied were randomly determined across the keystrokes (random phase), and the random phase was continued over 40 keystrokes. The random phase was used to prevent savings that are defined as faster relearning under a known environment. At 3 to 5 keystrokes after the random phase (the number of keystrokes was determined randomly), the artificial force was removed. Then, during the subsequent 40 keystrokes, the haptic device applied the artificial force to the finger in the direction of the X axis (adaptation phase). The artificial force direction was randomly chosen but was consistent in the adaptation phase. Because each pianist performed this task twice per tempo condition (i.e., 10 trials in total), the abduction and adduction directions of the artificial force during the adaptation phase were chosen once each per tempo condition and the order of the two direction in each tempo condition was randomly determined. Then, the artificial force was removed (washout phase), and the washout phase continued over 20 keystrokes.

We calculated the deviation of the fingertip force from the target force on the X-axis as an error index. We fitted the time course of the force deviation in the adaptation phase with a single state-space model:

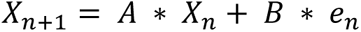

where X represents motor outputs at the n^th^ keystroke, e represents the error between the target and motor outputs at the n^th^ keystroke, A is the retention gain that represents how much motor outputs at a certain keystroke are utilized in motor outputs at its subsequent keystroke, and B is the feedback-related gain that represents how much motor outputs at a certain keystroke are influenced by the error obtained from the previous keystroke.

### Experiment 4-2

The purpose of this experiment was to test whether the effect of AHT on the target keystroke skill depended on the feedback-related gain. Thirty pianists participated in this experiment. They were categorized into either the FB group (n=15) or the no-FB (n=15) group. Before and after performing AHT, participants performed the target keystroke task at 5 different tempi (0.25, 0.5, 1, 2, and 3 Hz).

### Target keystroke task

The participants performed the target keystroke task with different tempi before and after AHT. In this task, the participants repetitively struck a piano key 20 times at a rate of 3, 2, 1, 0.5, 0.25 Hz using their right index fingers. They were instructed to produce 50% of the MVK. During the first 10 keystrokes, the participants received feedback information on the peak key-descending velocity value of each keystroke; this information was not provided during the subsequent 10 keystrokes. We calculated a coefficient of variation for the peak velocity values across the last 10 keystrokes as an index of the force production consistency. We repeated this task over 5 trials and calculated an average value of this index across the trials.

### Statistical analysis

The data were analyzed using a linear mixed effects model (LME) implemented in the lmerTest (*54*) package in R (https://www.r-project.org). Our model included the factors appropriate for the data among the group, time, tempo, and force level as fixed effects along with their interactions. Individual variability was assessed by including participants and participant*within factor interactions (when the number of within factors is 2 or more) as random factors. To determine the significance of the fixed effects, Type II Wald chi-squared tests were peformed using the Anova function in R with the car package (*55*). Multiple comparisons were performed using the emmeans package (*56*) in R. Degrees of freedom of such comparisons were calculated using the Kenward-Roger method, while p-values were adjusted using the Tukey method.

## Acknowledgments and funding sources

The present study was supported by JSPS Grant-in-Aid for Transformative Research Areas B (20H04093) for M.H, JST CREST (JPMJCR17A3, JPMJCR20D4) and JSPS Grant-in-Aid for

Transformative Research Areas B (20H05713) for S.F.

